# Convergent changes in gene expression associated with repeated transitions between hummingbird and bee pollinated flowers

**DOI:** 10.1101/706127

**Authors:** Martha L. Serrano-Serrano, Anna Marcionetti, Mathieu Perret, Nicolas Salamin

**Affiliations:** Department of Computational Biology, University of Lausanne, 1015 Lausanne, Switzerland; Conservatoire et Jardin botaniques de la Ville de Genève and Department of Botany and Plant Biology, University of Geneva, Chemin de l’Impératrice 1, 1292 Chambésy, Geneva, Switzerland

**Author notes:** these two authors contributed equally to the work (Martha L. Serrano-Serrano, Anna Marcionetti). Author for Correspondence: Nicolas Salamin, Department of Computational Biology, University of Lausanne, 1015 Lausanne, Switzerland, 0041 21 692 4154.

**Keywords:** Gesneriaceae transcriptomes, Neotropics, Flower development, Parallel evolution, Pollinator-mediated selection

## Abstract

The repeated evolution of convergent floral shapes and colors in angiosperms has been largely interpreted as the response to pollinator-mediated selection to maximize attraction and efficiency of specific groups of pollinators. The genetic mechanisms contributing to certain flower traits have been studied in detail for model system species, but the extent by which flowers are free to vary and how predictable are the genetic changes underlying flower adaptation to pollinator shifts still remain largely unknown.

Here, we aimed at detecting the genetic basis of the repeated evolution of flower phenotypes associated with pollinator shifts. We assembled and compared *de novo* transcriptomes of three phylogenetic independent pairs of Gesneriaceae species, each with contrasting flower phenotype adapted to either bee or hummingbird pollination. We assembled and analyzed a total of 14,059 genes and we showed that changes in expression in 550 of them was associated with the pollination syndromes. Among those, we observed genes with function linked to floral color, scent, shape and symmetry, as well as nectar composition. These genes represent candidates genes involved in the build-up of the convergent floral phenotypes.

This study provides the first insights into the molecular mechanisms underlying the repeated evolution of pollination syndromes. Although the presence of additional lineage-specific responses cannot be excluded, these results suggest that the convergent evolution of genes expression is involved in the convergent build-up of the pollination syndromes. Future studies aiming to directly manipulate certain genes will integrate our knowledge on the key genes for floral transitions and the pace of floral evolution.

**Data availability:** Raw Illumina reads will be available in the Sequence Read Archive (SRA) in NCBI database. The assembled transcriptomes and their annotation will by available in DRYAD repository. Details and accession ID will be provided at the time of the manuscript acceptance.

## Introduction

Cases of repeated evolution have attracted a large interest in evolutionary biology because they provide a replicated system to test long-standing questions in evolution. One of these questions is to which extent the independent origin of similar phenotypes is governed by shared molecular mechanisms (Steiner et al., 2009; Christin et al., 2010; Martin & Orgogozo, 2013). In plants, the origin and diversification of flower morphologies (Glover, 2014; Sauquet et al., 2017) and the occurrence of similar flowers in unrelated angiosperm species are striking examples of repeated phenotypic evolution (Thomson & Wilson, 2008; Ng & Smith, 2016). The genetic basis of these repeated changes is still largely unknown and identifying the mechanisms involved would greatly improve our understanding of the evolutionary success of angiosperms.

The widespread convergence in flower morphologies between angiosperm lineages is mainly explained by pollinator-driven selection that led to concerted evolution of a set of floral traits adapted to specific groups of pollinators (i.e. “pollination syndrome”, Fenster et al., 2004; Rosas-Guerrero et al., 2014). Our current knowledge of the genetic controls defining the pollination syndromes comes from quantitative genetic studies and experimental approaches on model species like *Petunia, Mimulus* and *Antirrhinum* (Stuurman et al., 2004; Galliot et al., 2006; Yuan et al. 2013). Outside these model systems, the trait that received most of the attention is the flower coloration. For example, the repeated evolution of red flower in hummingbird pollinated *Penstemon, Ipomoeae* and *Iochroma* is associated with concerted changes on gene regulation or on loss of function mutations inactivating specific genes (Des Marais & Rausher, 2010; Smith & Rausher, 2011; Wessinger & Rausher, 2015). However, the transitions between pollination syndromes involve a much larger suite of phenotypic traits that go beyond the coloration of flowers. While some of these floral traits may rely on a few genetic changes, others may have more complex genetic architecture. The transitions in pollination syndromes could, therefore, be much more complex than initially thought, and potentially controlled by different genetic solutions (Woźniak & Sicard, 2017).

Investigating the genetic basis of such a large array of floral traits in non-model species is difficult using traditional approaches. However, the replicated nature of these changes, coupled with the analyses of complete genomes or transcriptomes, provide new avenues to test for the genetic signatures and adaptive changes occurring during the convergent evolution of similar flower phenotypes (Clare et al., 2013). Many instances of repeated evolution occur during adaptive radiations, which involve large groups of mostly non-model organisms (Elmer & Meyer, 2011). These cases have received much less attention in the literature because functional experiments in these systems are hard to achieve. Further, the complexity of the phenotype targeted and the scarce genetic resources for many lineages have been limiting the possibilities of research in these groups (Ord & Summers, 2015). Fortunately, the advent of next-generation sequencing technologies enables the investigation of the genomic basis of repeated evolution in a variety of organisms (Lipinska et al., 2019). For example, the comparative transcriptomic and genomic study of the repeated evolution of eusociality in insects (Berens et al., 2015; Morandin et al., 2016) and bioluminescence in squids (Pankey et al., 2014) evidenced the presence of functionally shared elements and similarities in gene expression associated with these phenotypic convergences.

In this study, we aimed at detecting the genetic basis of the repeated evolution of flower phenotypes associated with pollinator shifts. Using a transcriptome-wide analysis, we quantified the differences in gene expression between three pairs of Gesneriaceae species, each corresponding to independent transitions between bee and hummingbird pollination systems (Fig. 1). So far, few studies have investigated the genetic controls of pollination syndrome morphologies in the Gesneriaceae family. Alexandre *et al.* (2015) surveyed second-generation population of hybrids between two *Rhytidophyllum* species and found that color and nectar differences between pollination syndromes were each controlled by a single QTL, but few candidate genes were identified. The high phenotypic convergence (Fig. 1B) and the lability of the plants-pollinators relationships in the Gesneriaceae (Fig. 1A), makes this family particularly suitable to study adaptive convergent evolution (Perret et al., 2007; Marten-Rodriguez et al., 2010; Clark et al., 2015; Serrano-Serrano et al., 2015; Serrano-Serrano et al., 2017a). The recent development of genomic resources in Gesneriaceae opens new possibilities to further explore the molecular mechanisms related to flower diversification in this lineage of tropical plants (Roberts & Roalson, 2017; Serrano-Serrano et al., 2017b). Here, we investigate to which extent the genetic basis of the bee and hummingbird floral specialization is primarily governed by shared changes in gene expression. Our results allow us to quantify the extent of convergent responses, and to identify candidate genes and pathways that are associated with the repeated evolution of flower morphological changes underlying pollinator specificity.

**Figure 1.**
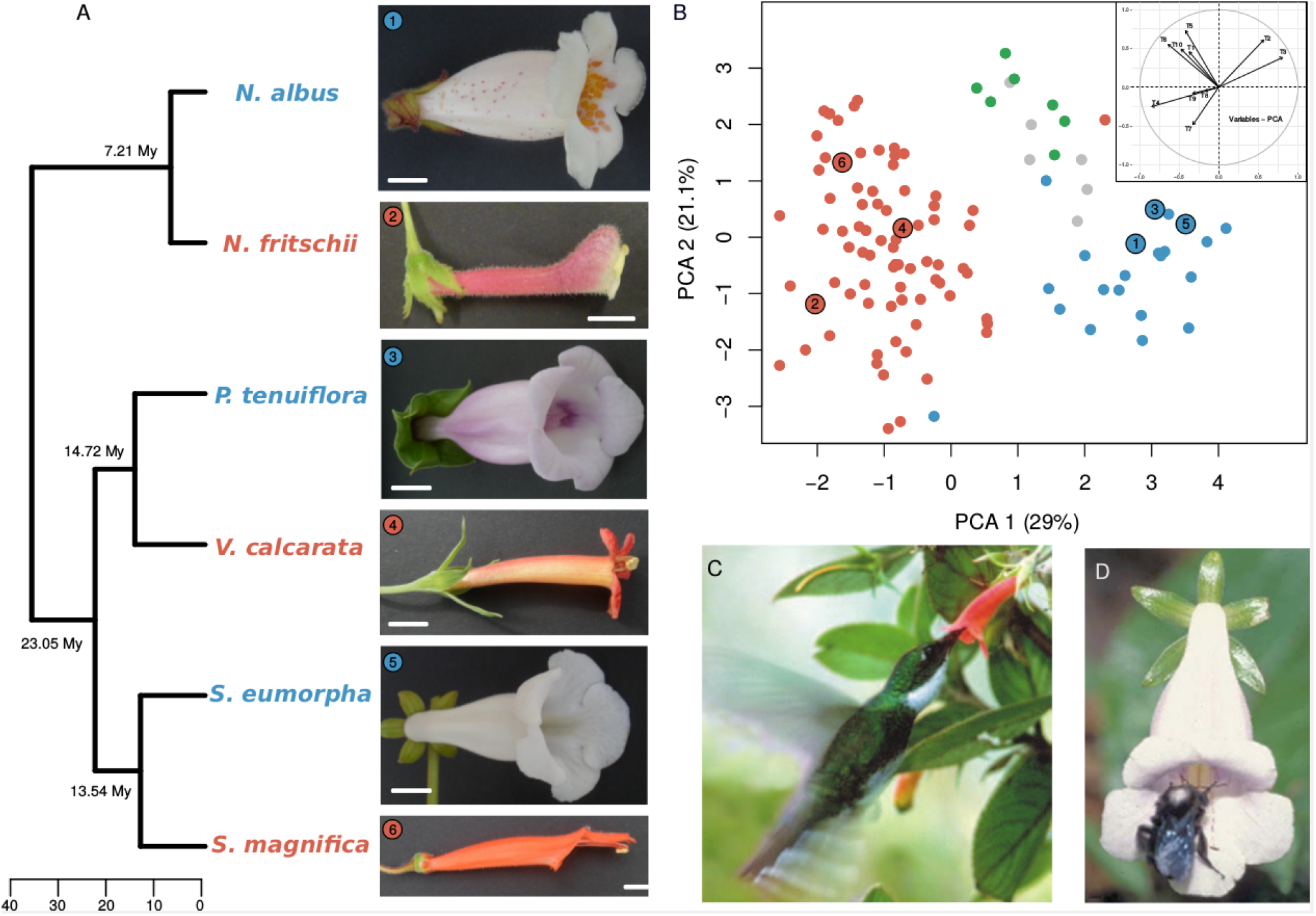
Experimental setting. *A)* Phylogenetic relationships of the six Gesneriaceae species and their morphology. White bar represents 1 cm. *B)* Principal component analysis for ten floral measurements in all the species within the genera *Nemantanthus* and *Sinningia* (see Table S2). Colors correspond to the pollination syndromes (blue= insect, red= hummingbird, green= bat, see Serrano-Serrano *et al.* 2017b). Numbers correspond to the species 1= *N. albus*, 2= *N. fritschii*, 3= *P. tenuiflora*, 4= *V. calcarata*, 5= *S. eumorpha*, 6= *S. magnifica. C) V. calcarata* visited by *Leucochloris albicolis* (photo from SanMartin-Gajardo & Sazima, 2005). *D) S. eumorpha* visited by *Bombus morio* (photo from SanMartin-Gajardo & Sazima, 2004).

## Results

We selected three phylogenetic independent pairs of related species that differ in floral phenotype and functional groups of pollinators (Table S1). These six species show two contrasting pollination syndromes: bee-pollinated (*Nematanthus albus, Sinningia eumorpha*, and *Paliavana tenuiflora*) and hummingbird-pollinated (*N. fritschii, S. magnifica*, and *Vanhouttea calcarata*, see Fig. 1A). We investigated similarities in their floral morphologies and found that floral traits defining their morphospace (Table S2; data from Perret et al., 2007, Serrano-Serrano et al., 2015; see Methods) showed a clustering of species according to their pollinators (Fig. 1B). Flower traits related to these distinct groups of pollinators are mostly associated with floral shape (bell-shape versus tubular corollas), length of the anther/stigma (short versus exerted), and color with mostly whitish versus red, pink or orange corollas (Fig. 1B and inset). These results and the phylogenetic independence of the species pairs that we selected demonstrate the convergent nature of the bee and hummingbird pollination syndromes among the sampled species (Fig. 1A), and across the multiple pollinator switches in the Gesnerioidea subfamily (Serrano-Serrano et al., 2017a). The selected species are therefore particularly suitable to investigate the genetic bases of the repeated evolution of flower phenotypes associated with pollinators shifts, and thus for the study of adaptive convergent evolution.

### *De novo* transcriptomes assembly, annotation and orthology inference

For each species, we sequenced the RNA extracted from the leaf and floral tissues at three different developmental stages, we assembled *de novo* the transcriptomes, annotated them and checked their quality (see Material and Methods, SI S1). The transcriptomes obtained for the four species (*P. tenuiflora, V. calcarata, N. albus*, and *N. fritschii*) were added to the data of *S. eumorpha* and *S. magnifica* from Serrano-Serrano *et al.* (2017b). The final number of inferred proteins per species ranged from 136,772 to 171,808 (Table S4). Additional information on the quality and annotation of the transcriptomes is provided in Supplementary Information S1.

Orthology inference from the predicted protein of the six species resulted in the identification of a total of 96,429 orthologous groups (OG), with most of them (36,201) not further considered because they are composed of proteins found in a single species (Figure S3A). We were interested in investigating changes in gene expression potentially associated with the convergent evolution of pollination syndromes and we therefore focused on OGs composed by species enabling the contrast of the two pollination syndromes. This resulted in a total of 14,059 OGs composed by 3 (“pollination-syndorme-specific OG”), 4 (“four-species OGs”), 5 (“five-species OG”) or 6 species (“1-to-1 OG”), which were analyzed for differential expression linked with pollination syndromes (see SI S2 for more information).

### Overall comparison of the expression profile of the sequenced libraries

Gene expression for each library and species was quantified by mapping the reads to the corresponding assembled transcriptome, quantifying the number of mapped reads and normalizing these counts (see Material and Methods, SI S3 and S4). A PCA on the normalized expression level of the 7,287 1-to-1 OGs showed that the global expression profiles primarily clustered RNA-seq libraries by tissues (floral vs leaf tissues, PC1= 16.79% variance) and lineages (PC2= 14.02% variance). The next two axes of variation separated the floral developmental stages (small buds, middle buds, adult flower), with PC3 and PC4 axes explaining jointly an additional 19.13% of the variation (see Fig. 2).

**Figure 2.**
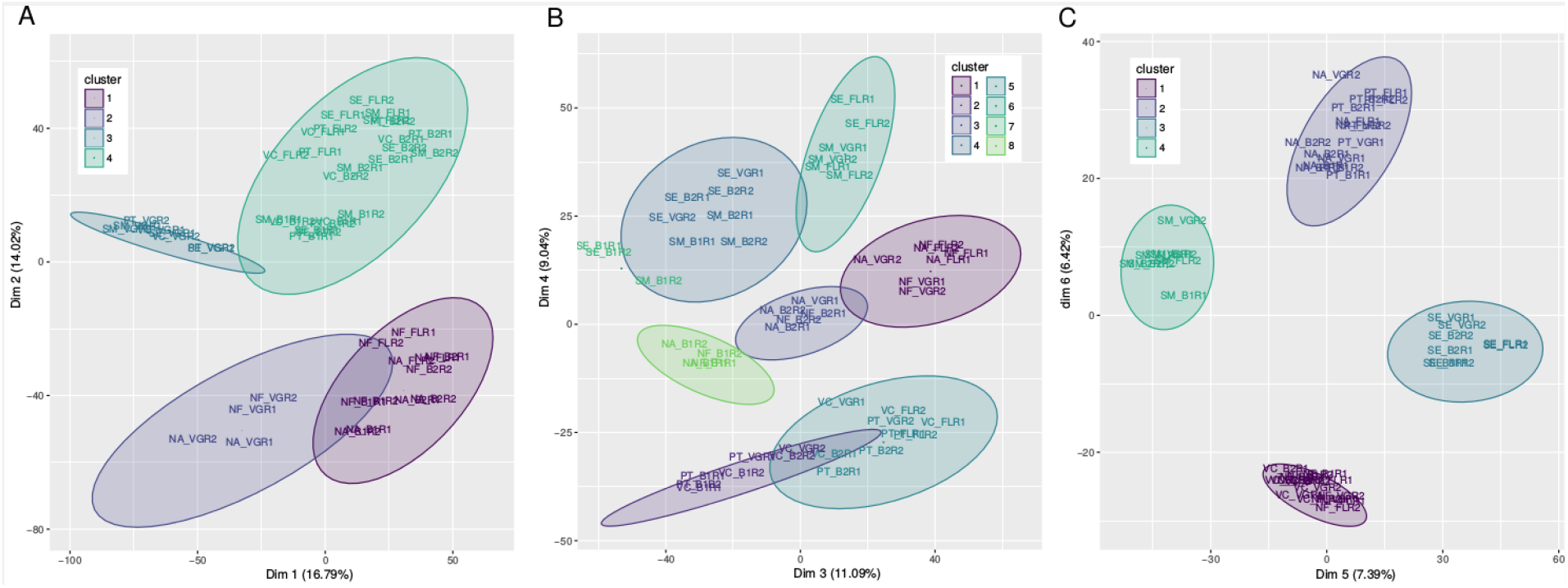
Principal Component Analysis of the normalized RNA expression levels for all species. The proportion of variance explained by the PCA is indicated in parenthesis on each axis. Panels A, B, and *C* represent the first six axes. Colors determine the number of clusters using k-means. Labels on the libraries indicate the species (i.e. SE, *S. eumorpha*), the developmental stage (i.e. B1, early bud) and, the replicate number (i.e. R1, replicate 1). Information on the cluster composition is provided in Table S7.

Although most of the variance in gene expression was explained by tissue, stage and species, a separation driven by the pollination type was also observed in axes PC5 and PC6, which together explained an additional 13.81% of the variance. This result suggests that species with similar pollination syndromes may display similar expression patterns in some of the analyzed genes, but that it is not the major source of variation found in the transcriptome data.

### Differential gene expression analysis between pollination syndromes: 1-to-1 OGs

For 1-to-1 OGs, two different approaches were employed to investigate the differentially expressed genes (DEGs) between bee- and hummingbird-pollinated species. First, in the “pairwise” differential expression analysis, we searched for DEGs at each floral developmental stage in each pairs of related species. We then compared the pairwise analyses to investigate whether the DEGs were shared between species-pairs (Figure 3A-C). Second, in the “global” differential expression analysis, we directly contrasted all bee-pollinated against all hummingbird-pollinated species, thus considering species with the same pollination syndromes as replicates (Figure 3D). While both approaches can detect the clearest cases of concerted changes in expression linked with the pollination syndromes, for less obvious cases the two approaches are complementary (see SI S5 and Figure S6).

**Figure 3.**
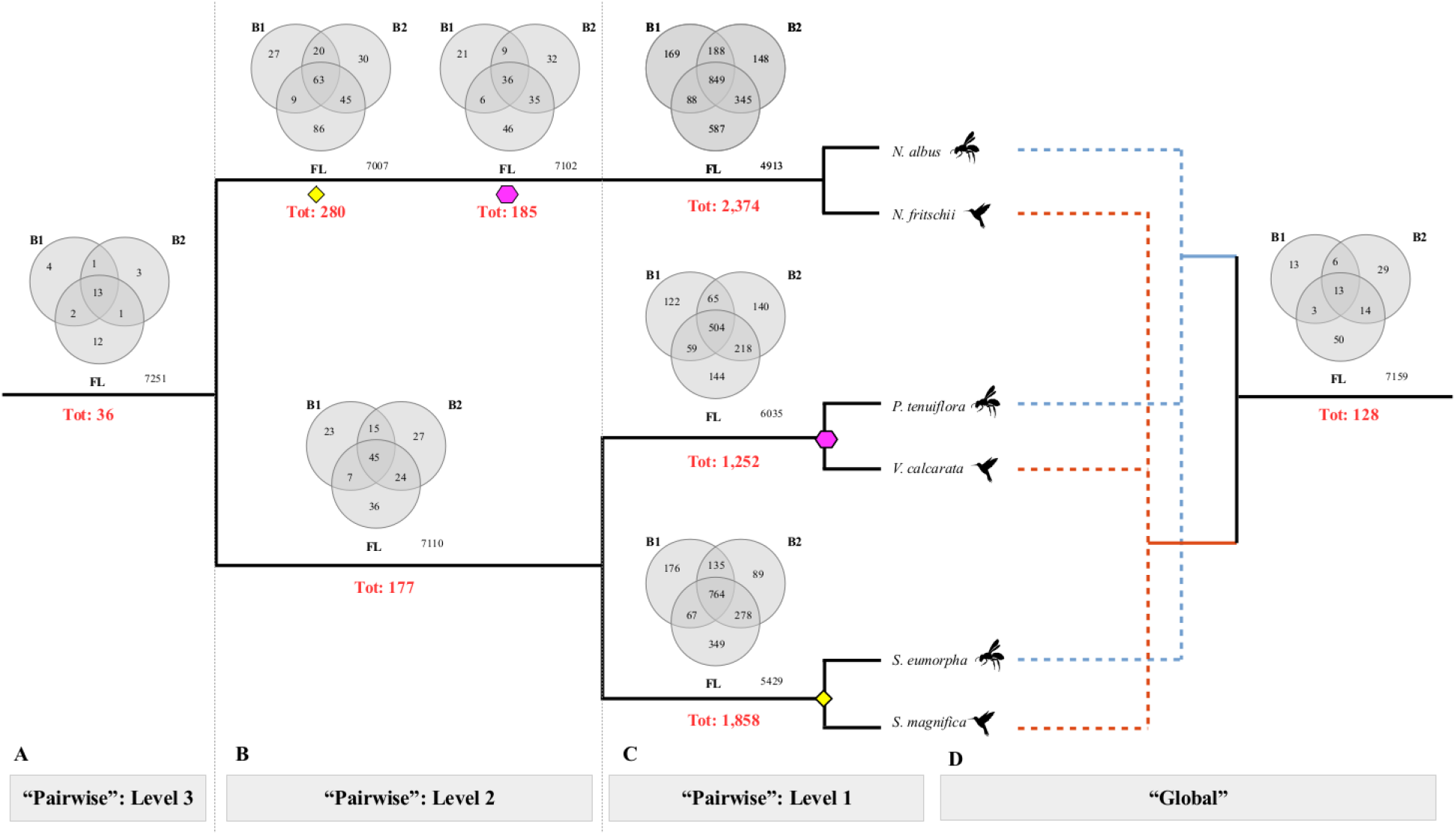
Schematic phylogenetic tree of the six Gesneriaceae species and the differential expression analyses performed. **A, B, C** panels represent the “pairwise” analysis, with genes found differentially expressed in pairwise species comparison (Level 1, **C**), found shared between two species pairs (Level 2, **B**) and shared between the three pairs (Level 3: **A**). Panel **D** represents results for the contrast performed in the “global” analysis. For all panels, the number of differentially expressed genes between floral phenotypes is displayed in each branch for each comparison. For each comparison, the total number of DEG is reported below the branch in red. The different expectation for the results of the two analyses are reported in SI S5 and Figure S6.

#### Pairwise differential expression

Out of the 7,287 1-to-1 OGs analyzed, 2,374, 1,252 and 1,858 DEGs were found between, respectively, *N. albus* vs *N. fritschii, P. tenuiflora* vs *V. calcarata* and *S. eumorpha* vs *S. magnifica* (Figure 3C). For all species pairs, around 60% of the DEGs were functionally annotated (see Tables S8-S10 for a complete list). When considering the total number of DEGs in each developmental stage separately (B1, B2 and FL), we observed a trend of increasing number of DEGs with the progressing floral development (*Nematanthus* pair: B1 = 1,294, B2 = 1,530, FL = 1,869; *Sinningia* pair: B1 = 1,141, B2 = 1,266, FL =1,458; *Paliavana-Vanhouttea* pair: B1 = 750, B2 = 927, FL = 925). In all species-pairs comparison, the number of DEGs with higher expression in bee-pollinated species (arbitrarily set to category “1”) was similar to the number of DEGs with higher expression in hummingbird-pollinated species (category “-1”; *Nematanthus* pair: 1,107 and 1,233 DEGs for respectively “1” and “-1” categories; *Paliavana-Vanhouttea* pair: 641 and 599 DEGs for respectively “1” and “-1” categories; *Sinningia* pair: 933 and 910 DEGs for respectively “1” and “1” categories; Figure S7).

The number of DEGs shared betwee two species-pairs considerably dropped with a total of 280, 185 and 177 DEGs found to be shared between, respectively, the *Nematanthus* and *Sinningia* pairs, the *Nematanthus* and *Paliavana-Vanhouttea* pairs, and the *Sinningia* and *Paliavana-Vanhouttea* pairs (Figure 3B, Table S11). This number further decreased when considering DEGs shared between the three species pairs with only 36 being consistently differentially expressed between all bee- and hummingbird-pollinated species (Figure 3A, Table 1A, Table S12). Around 70% of the 36 DEGs shared between the three species-pairs were functionally annotated (Table S12) and most of these DEGs were found only in the adult flower (12 genes) or were shared between the three developmental stages (13 genes, Figure 3A).

We investigated whether the overlap of DEG between species-pairs at each developmental stage were only due to random expectation (SI S8). We found that for most of the comparisons, the number of shared DEGs between two species-pairs (Figure 3B) and between all species pairs (Figure 3A) were higher than expected by chance (Figure S8), which confirms an association between the convergent pollination syndromes and the presence of shared DEGs.

**Table 1:**
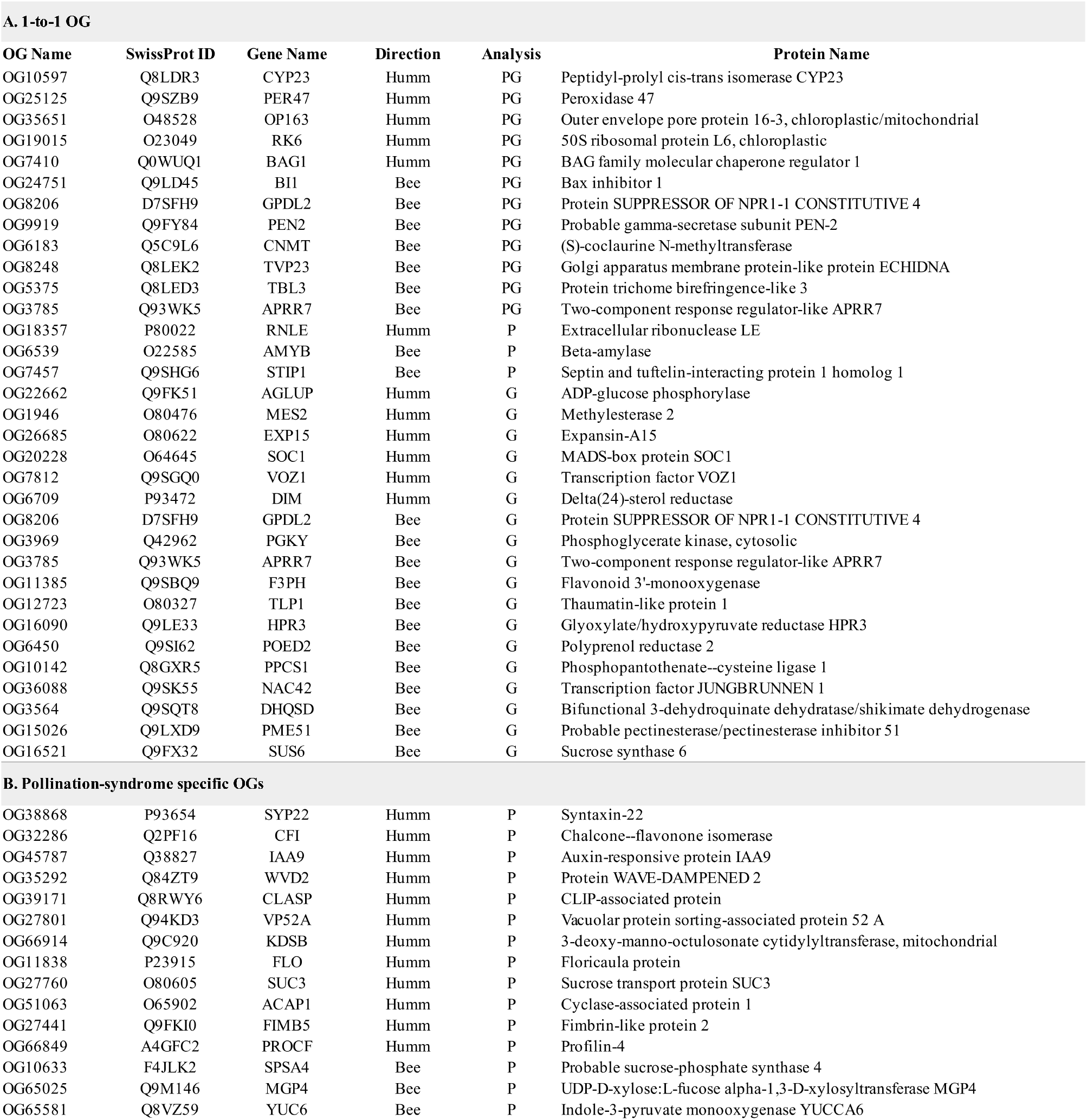
Partial list of differentially expressed genes showing concerted changes in gene expression associated to the pollination syndromes. A: 1-to-1 OG detected differentially expressed and linked with the pollination syndromes in the “pairwise” (P), “global” (G) or both (PG) analysis. B: Pollination-syndrome specific OGs with the expression linked with the two different pollination syndromes. The direction of the expression is reported in the “Direction” column, with Humm and Bee corresponding to higher expression in hummingbird- and bee-pollinated species, respectively. Complete list is available in the SI, Tables S12, S13 and S14

#### Global differential expression

Out of the 7,287 1-to-1 OGs analyzed, we found 128 DEGs between all bee- and hummingbird-pollinated species (Figure 3D). Around 70% of the DEGs were functionally annotated (see complete list in Table S13). The highest number of DEGs was found for the adult flower (50 DEGs) and we observed a similar trend as for the pairwise analysis of an increase in the number of DEGs with floral development (35, 62 and 80 DEGs respectively for B1, B2 and FL, Tables S13). The number of DEGs with higher expression in bee-pollinated species (i.e. 67 genes) was similar to the number of DEGs with higher expression in hummingbird-pollinated species (i.e. 61 genes).

#### Comparison of the results from “pairwise” and “global” analyses

The number of 1-to-1 OGs with concerted expression changes associated with the pollination syndromes is different depending on the considered analysis (36 DEGs for “pairwise”, 128 for “global”). The presence of DEGs specific to the two analyses was expected as the two approaches are complementary (SI S5, Figure S6).

A total of 29 DEGs are shared between the two analysis. The shared DEGs show a clear pattern of differential expression linked with the pollination syndromes (see Fig. 4A for examples). Differences in expression between pollination types were also observed for genes identified with the “pairwise” approach, although there is an absence of absolute difference in expression between bee- and hummingbird-pollinated species (Figure 4B). Finally, the expression of DEGs specific to the “global” analysis is shown in Figure 4C, with genes showing lower variation in expression between pollination syndromes (OG2262) and/or high variation within replicates in one or more species-pairs (OG36088, OG20228). These characteristics prevented those genes to be detected as differentially expressed by the “pairwise” analyses, although their expression patterns are associated with the pollination syndromes.

**Figure 4.**
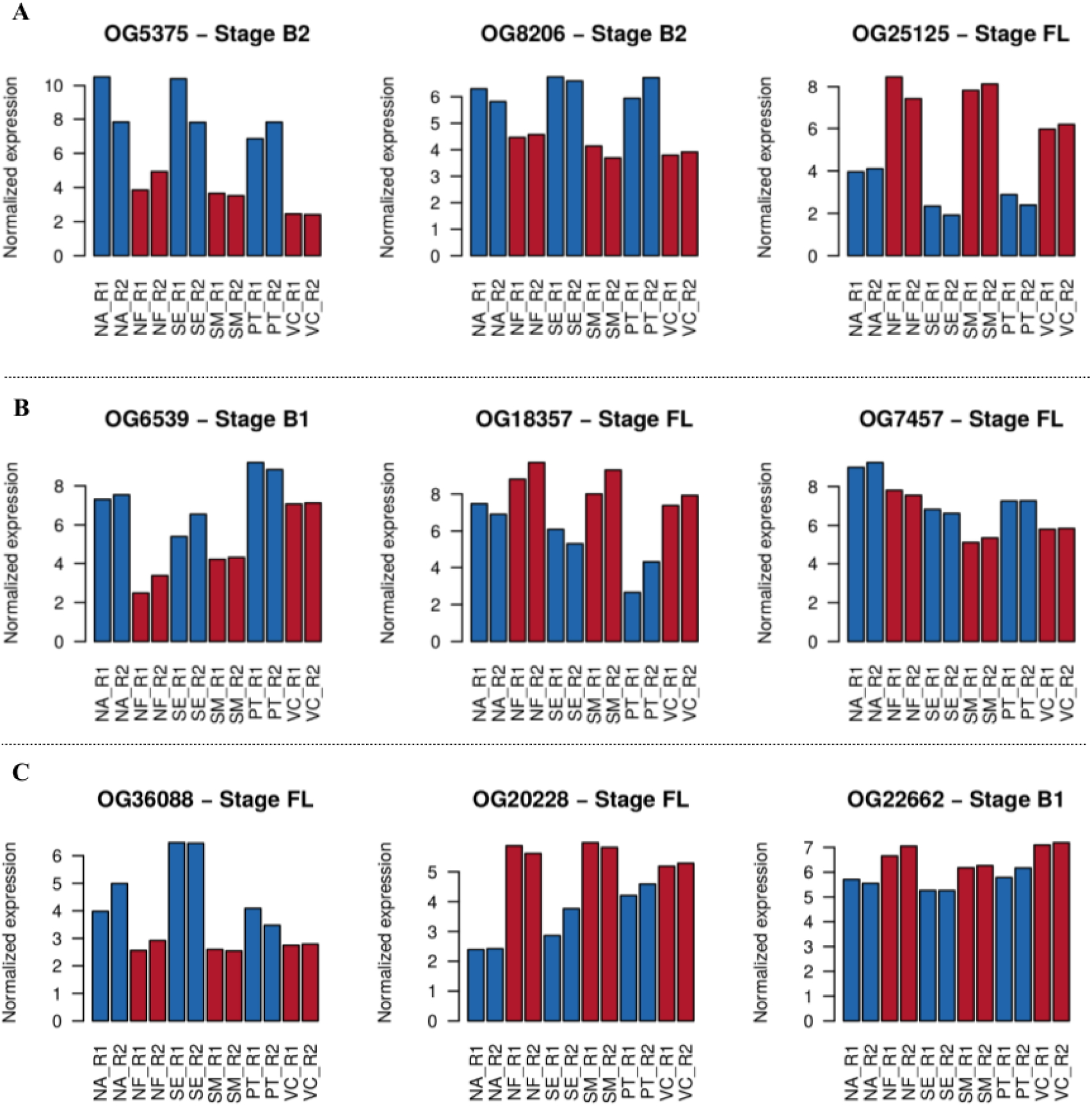
Examples of 1-to-1 OG found concordantly differentially expressed between pollination syndromes. OGs detected with both “pairwise” and “global” analysis are shown in panel **A**, while panel **B** and **C** show the expression of OGs detected respectively with “pairwise” and “global” analysis. The expression is shown as the log2(Normalized Expression). NA, NF, SE, SM, PT and VC correspond respectively to *N. albus*, *N. fritschi, S. eumorpha*, *S. magnifica*, *P. tenuiflora* and *V. calcarata*. R1 and R2 correspond to the two replicates. The developmental stage (B1, B2 or FL) having the represented expression is reported in the title of each graph.

### Differential gene expression analysis between pollination syndromes: pollination-syndrome-specific OGs, four- and five- species OGs

We investigated the changes in gene expression potentially associated with the convergent evolution of pollination syndromes by also analyzing whether the expression of the pollination-syndrome-specific, four-species and five-species OGs was associated with the phenotypes.

Out of a total of 253 OGs specific to bee-pollinated species, 198 were found to be significantly expressed in the three species while absent in the hummingbird-pollinated species (see Material and Methods, Figure 5A). Similarly, out of the total of 313 OGs specific to hummingbird-pollinated species, 217 were found significantly expressed in the three species while absent in the bee-pollinated species (Figure 5B). Out of the 198 bee-pollinated species specific OG and 217 hummingbird-pollinated species specific OGs, 117 and 126 were functionally annotated (Table 1, Table S14).

**Figure 5.**
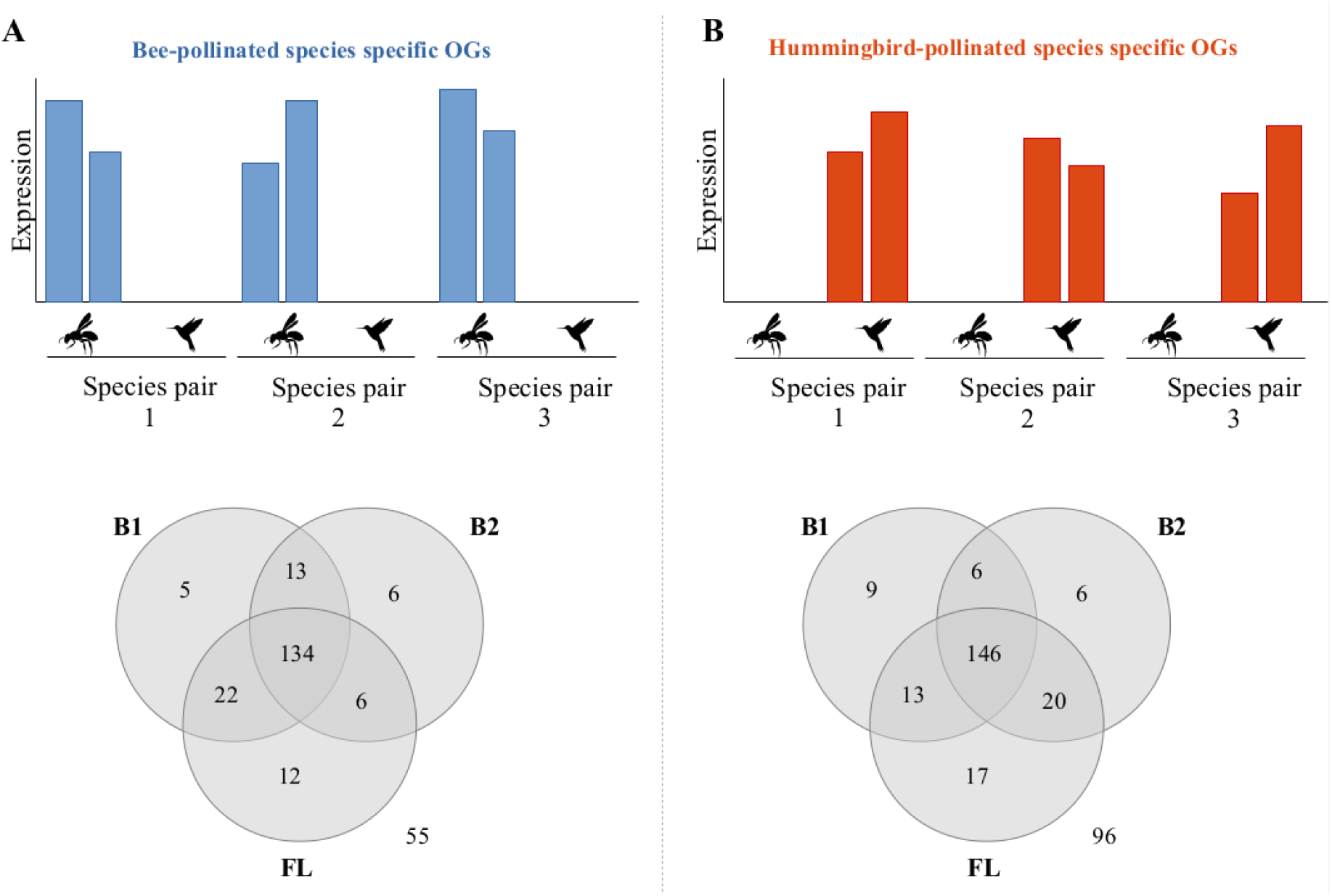
Schematic representation of pollination-syndrome-specific OG, with bee-pollinated species specific OG (A) and hummingbird-pollinated species specific OG (B). In the bottom part of the figure, the total number of genes showing a significantly higher expression in bee-pollinated species specific OG **(A)** and hummingbird-pollinated species specific OG **(B)** is reported.

We found 6,027 OGs present in four of the six species. Of these, 986 were composed by the three bee-pollinated species plus one hummingbird-pollinated species, while 1,043 were formed by the three hummingbird-pollinated species and one bee-pollinated species. Similarly, out of the 4,209 OGs composed by a total of five species, 2,140 were composed by the three bee-pollinated species plus two hummingbird-pollinated species, while 2,069 were formed by the three hummingbird-pollinated species and two bee-pollinated species. For both categories, we searched for genes with an expression pattern that was associated with the pollination syndromes (see Material and methods, SI S7), but no DEG was detected.

### Differential gene expression analysis between pollination syndromes: overall results

In total, we investigated whether the expression of 14,059 OGs was associated with the convergent pollination syndrome phenotypes. When considering all OGs categories and all analyses together, we found a total of 550 genes (415 pollination-syndrome specific OGs, 135 1-to-1 OGs considering both “pairwise” and “global” analysis) with an expression pattern significantly linked with the pollination syndromes. A functional annotation was available for 302 of them.

A protein-protein network reconstructed with STRING resulted in a network having significantly more interaction that expected (minimum required interaction score: 0.9, STRING PPI enrichment p-value: 0.00939; Figure S9). This suggests that the detected genes are at least partially connected through biologically relevant functions.

KEGG enrichment analysis resulted in the enrichment of the three wide KEGG pathways: metabolic pathways (ID: ath01100, fdr=2.5×10^−10^), biosynthesis of secondary metabolites (ID: ath01110, fdr=1.92×10^−06^) and biosynthesis of amino acids (ID: ath01230, fdr=4.6×10^−04^). GO enrichment analysis resulted in the enrichment of a total of 171 terms (Table S15). Within them, GO linked with cell growth (e.g. GO:0016049, GO:0009826, GO:0048588), pollen tube development (e.g. GO:0048868, GO:0009860), cytoskeleton organization (e.g. GO:0007010, GO:0000226), carbohydrate metabolic process (e.g. GO:0016052, GO:1901137, GO:0016051) and amino acid metabolic process (e.g. GO:0006520, GO:0008652) were present.

## Discussion

The mechanisms underlying the repeated evolution of similar phenotypes can involve modifications in homologous genes or in distinct enzymes and pathways (Christin et al., 2010). These differences may reflect the complexity of a phenotype or the level of genetic and molecular redundancy, which both translate in the potential of a phenotype for evolutionary change. Current knowledge of the genetic control of flower-pollinator specificity is largely derived from the study of model species from phylogenetic disparate plant lineages, such as *Petunia, Mimulus* and *Antirrhinum* (Yuan et al., 2013; Woźniak & Sicard, 2017). However, these systems, which all involve a single or few closely related species, are not appropriate to investigate the convergent nature of pollinator-driven flower evolution within plant radiations. Here, we identified the convergent changes in gene expression between bee- and hummingbird-pollinated flowers using three species pairs that belong to one single evolutionary radiation in the Gesnerioideae. This lineage is mainly distributed in the Neotropics and it has undergone multiple pollinator switches (Serrano-Serrano et al., 2017a). It represents thus a promising framework to examine the magnitude of repeated changes existing outside of model plant species (Chanderbali et al., 2016). Our experimental design was challenging from an evolutionary comparative transcriptomic perspective (e.g. the comparison of different developmental stages or the comparison of transcriptomic data between species, as reported in Roux et al., 2015), but it enabled us to characterize the repeated evolution of floral phenotypes in non-model systems by taking advantage of natural replication of evolutionary events.

Before considering the molecular similarities underlying the flower pollination syndromes, we discuss the global expression patterns. The overall characterization of the gene expression in the six species indicates that the expression profiles are similar between all of them, and the major distinctness between libraries occurs between tissues followed by stages of development (Fig. 2*A-B*). This differentiation in transcriptional programs has been shown in comparisons within and between species (Wellmer et al., 2006; Pazhamala et al., 2017), suggesting a deep conservation of gene expression patterns within tissues and developmental stages even at large evolutionary scales (Brawand et al., 2011).

### Function of genes showing convergent changes in expression between pollination syndromes

When investigating the genetic elements that were repeatedly recruited in similar phenotypes, we found that 3.9% (550 out of the 14,059) of the genes analyzed showed changes in expression associated with the pollination syndromes in at least one floral developmental stage. Some of these genes are likely to control floral traits associated with pollinator switches between hummingbird and insects.

The gene ontology (GO) enrichment analysis performed on the DEGs resulted in an enrichment of GO categories linked with cell growth and cytoskeleton organization. Although these terms are overall broad, they potentially include genes involved in the build up of the convergent floral shape and symmetry in the two pollination syndromes. Similarly, an enrichment in GO categories linked with the development and growth of pollen tube was also observed for the genes with parallel differential expression. This suggest the presence of genes involved in the differential elongation of pollen tube in the two different pollination syndromes (Fig. 1A).

The presence of GO enrichment terms such as developmental growth and morphogenesis suggest that, for some complex traits (such as the floral shape and symmetry), several genes with convergent differential expression may be required to acquire the convergent phenotype. The absence of enrichment for other interesting GO terms could be associated with simple traits that necessitate only one or few genes with a convergent differential expression to switch to the other phenotype.

Below, we discuss the functions of some candidate genes that may be involved in the convergent switches between pollination syndromes and could be associated with the different floral traits observed in each species pair.

### Floral color

One of the traits that has been the most widely studied in pollination syndromes and for which the molecular basis of natural variation is starting to emerge is the floral color (Sheehan et al., 2012). Although some variation in the coloration exists, the species selected in this study clearly displayed a switch in the floral color depending on the pollinator type, with essentially white flowers for bee-pollinated species, and reddish flowers in hummingbird-pollinated species (Fig. 1). Anthocyanins, a class of flavonoid, are the predominant floral pigment in angiosperms (Winkel-Shirley, 2001) and variation in either the coding sequence or the expression of enzymes of the flavonoid biosynthetic pathway are associated with change in floral color and switches in plant pollinators (reviewed in Sheehan et al., 2012).

We found three DEGs between bee- and hummingbird-pollinated species having known functions related to the production of anthocyanins. The first is OG11385, which was annotated as flavonoid 3’-monoxygenase (*F3’H* or *F3PH*) and is responsible for converting dihydrokaempferol into dihydroquercetin during the biosynthesis of anthocyanins. An increased expression of this gene potentially deviates the metabolic pathway towards non-red anthocyanins (Fig. 1 in Rausher, 2008) and other flavanones by substrate competition (Sharma et al., 2012; Yuan et al., 2016). In our species, the *F3’H* showed a significantly decrease expression in hummingbird-pollinated species (Fig. 4). A lower expression of *F3’H* in hummingbird-pollinated flowers compared with bee-pollinated has been previously reported for *Ipomoea* and *Iochroma* species (Des Marais et al., 2010; Smith et al., 2013), and it was usually associated with an inactivation (i.e. sequence degeneration and pseudogenization) of *F3’5’H* as a way to block permanently the access to non-red anthocyanins. This mechanism seems to underlie the evolution of red-flowers in *Penstemon* species, and it may constraint the evolutionary reversals of red flowers in that lineage (Wessinger & Rausher, 2014). Although the degenerative scenario for *F3’5’H* gene is unlikely in the Gesneriaceae because of the many reversals to bee-pollination (Serrano-Serrano et al., 2017a), the concerted decrease in *F3’H* expression in hummingbird-pollinated species may still be associated with the decrease in non-red anthocyanins and thus the maintenance of reddish flowers observed in these species. The presence of *F3’H* expression in bee-pollinated species despite the overall whitish color of the flowers may be explained by the presence of localized purple pigmentation (Fig. 1).

Another gene involved in the anthocyanin pathway found to be differentially expressed between bee- and hummingbird-pollinated species was the OG32286, which was annotated as the chalcone-flavonone isomerase (CHI). This enzyme is involved in the flavonoid biosynthesis, upstream from the anthocyanin biosynthesis pathway (Fig. 1 in Rausher, 2008). Suppression of *CHI* in transgenic tobacco plants resulted in flavonoid components and transition of floral color from red-purple to whitish (Nishihara et al., 2005). We detected the expression of *CHI* only in hummingbird-pollinated species. In bee-pollinated species this gene was either not expressed or expressed at very low levels, potentially leading to the overall white color of flower in bee-pollinated species (Fig. 1).

Finally, the last gene found to be differentially expressed between bee- and hummigbird-pollinated species was the OG36088, which was annotated as the transcription factor JUNGBRUNNEN 1 (*JUB1*). This gene showed a significantly increased expression in bee-pollinated species. It has been previously observed in *Arabidopsis thaliana* overexpressing *JUB1* that the expression of important enzymes of the anthocyanin pathway was significantly down-regulated (Wu et al., 2012). Taken together, these results suggests that the convergent evolution of floral color in Gesneriaceae is potentially due to the convergent changes in expression of genes from the flavonoid biosynthetic pathway.

### Nectar composition

Nectar plays a key role in plant reproduction as it reward pollinators (Proctor et al., 1996). Nectar is mainly composed by three sugars: the disaccharide sucrose and the hexose monosaccharides fructose and glucose (Percival, 1965). The ratio of sucrose over fructose and glucose (i.e. nectar sugar composition) has often been linked with different classes of pollinators. For instance, flowers pollinated by hummingbirds, butterfly, moths and long-tongues bees show a sucrose-rich nectar, while hexose-rich nectar has been found in flowers pollinated by short-tong bees and flies (Baker & Backer, 1983, 1990; Freeman et al., 1984; Lammers & Freeman, 1986; Elisens and Freeman, 1988; Stiles & Freeman, 1993; Backer et al., 1998; Dupont et al., 2004). In the *Sinningia* tribe, the sucrose was shown to be the dominant constituent in the nectar of both hummingbird- and bee-pollinated flowers, with however a trend for higher sucrose/hexose ratios in hummingbird-pollinates species (Perret et al., 2001).

A gene related to sucrose was found differentially expressed between the pollination syndromes and may be potentially linked with differences in sucrose composition in flowers. This gene was the Sucrose synthase 6 (*SUS6*, OG16521), an enzyme which cleaves sucrose, resulting in the production of glucose and fructose. This gene was more highly expressed in bee-pollinated species, which correlates with the reduced sucrose/hexose ratio trend observed in Perret et al. (2001). A similar expression pattern of this gene was observed in the nectaries of *Arabidopsis thaliana* ecotypes, where a low sucrose to hexose ratio was linked to increased expression of sucrose synthases (Fallahi et al., 2008). An increased activity of the sucrose synthase may thus reduce the accumulation of sucrose in the nectaries, resulting in a higher hexose concentration (Fallahi et al., 2008).

### Floral scent

Floral scent is obtained by the combination of different volatiles mostly belonging to fatty acid derivatives, terpenoids, or benzenoids (Sheehan et al., 2012). The number and concentration of compounds provide species-specific olfactory cues, and different pollinator animals (such as bats, bees and hummingbird) are able to sense floral scent and disentangle among odor differences (von Helversen et al., 2000; Wright et al., 2002; Kessler et al., 2008; Knudsen et al., 2004). Differences in the production of volatiles compounds is therefore expected when comparing bee- and hummingbird-pollinated species.

The gene OG15474 (Phenylalanine N-monooxygenase, *CYP79A2*) showed an increased expression in the flowers from bee-pollinated species. This protein converts the L-phenylalanine to phenylacetaldoxime, a precursor of benzylglucosinolate (Wittstock & Halkier, 2000), and was found to be a candidate scent gene in *Brassica rapa* (Cai et al., 2016). Another gene, the OG66797 was found to be expressed in hummingbird-pollinated species and was annotated as 5-enolpyruvylshikimate-3-phosphate synthase (*EPSPS*). This protein is part of the Shikimate pathway and is producing the precursors necessary for the production of volatile compounds in scented *Petunia* (Sheehan et al., 2012).

### Floral size, shape and symmetry

Other traits associated with convergence depending on the pollinator type are the floral shape and symmetry. Indeed, bee-pollinated species have bell-shaped corollas, while hummingbird-pollinated species show tubular morphologies (Fig. 1A).

Genes found differentially expressed between pollination syndromes includes both genes with functions linked to cell elongation and cell shape. For instance, the OG6709 was annotated as Delta(24)-sterol reductase (or Diminuto). This protein plays a critical role in the process of plant cell elongation (Takahashi et al., 1995; Schultz et al., 2001). Similarly, the OG26685 was annotated as Expansin-A15 (*EXP15*), a protein that causes loosening and extension of plant cell walls (Zenoni et al., 2011). Two other genes showed functions associated with cell expansion and cell polarity: *CLASP* (CLIP-associated protein, OG39171; Kirik et al., 2007) and *WVD2* (Protein WAVE-DAMPENED 2, OG35292; Yuen et al., 2003). All these genes were more highly expressed in hummingbird-pollinated species compared to bee-pollinated species, suggesting a potential role in the cell elongation of hummingbird-pollinated flowers, which could lead to the tubular morphology.

### Lineage-specific expression linked with different pollination syndromes

In this study, we found around 4% of the genes that show parallel changes in expression associated with the convergent acquisition of pollination-syndrome. Our results suggest that the convergent evolution of genes expression is involved in the build-up of the specific pollination syndromes. However, this does exclude the presence of lineage-specific responses. Indeed, some genes associated with pollination syndromes in other organisms were found differentially expressed in only some of the species pairs. Examples are the dihydroflavonol-4-reductase (*DFR*), chorismate mutase 1 (*CM1*), RADIALIS (*RAD*) and bidirectional sugar transported SWEET9 (*SWEET9*) genes, which we discussed in more details.

The gene *DFR* (OG12343) is involved in antocyanin biosynthesis and it showed a significantly higher expression only in *S. mangnifica* compared to *S. eurmopha* (Table S10). The expression of this gene may therefore be involved in a further differences in coloration between the two *Sinningia* species. The gene *CM1* (OG29505) only showed a significantly higher expression in *N. albus* compared to *N. fritshcii* (Table S8). This gene is involved in the production of floral volatiles in *Petunia* (Colquhoun et al., 2010) and thus may play a role in generating differences in scent between the *Nemathantus* species. The gene *RAD* (OG35772) showed a higher expression in *N. fritschii* compared to *N. albus* (Table S8), and it is a gene involved in the dorsoventral asymmetry of flowers (Corley et al., 2010). Finally, the *SWEET9* gene (OG23655) showed an increased expression in S. *eumorpha* and *P. tenuiflora* compared to *S. magnifica* and *V. calcarata.* This gene is involved in the production of nectar (Roy et al., 2017) and thus may be associated with differences in nectar production between these species, but not in the *Nemathantus* genus. While the lineage-specific expression of these genes may represent alternative routes to the build-up of the convergent pollination-syndromes phenotypes, the differences in expression of these genes may also be tuning some subtle differences in phenotypes when comparing the same pollination syndromes.

## Conclusion

Here, we developed an extensive transcriptomic resource for the Gesneriaceae family that, together with the one presented in Serrano-Serrano et al. (2017b), provides a comprehensive dataset for comparative analyses of this species-rich Neotropical plant group. The large diversity of pollination forms found in this group enabled us to take advantage of the repeated evolution of floral phenotypes to investigate the genetic basis of complex but key traits associated with the evolution of pollination syndromes.

By comparing the gene expression in bee- and hummingbird-pollinated flower, we found that around 4% of the genes showed parallel changes in expression associated with the convergent acquisition of pollination-syndromes. Among those genes, we identified several that have functions associated with reported differences in the floral phenotype of the two pollination syndromes (i.e. floral color, scent, shape and symmetry, as well as nectar composition). However, our study further gives new insights into the genetic make-up of pollinator transitions by providing new candidate genes involved in one of the most essential evolutionary process linked with plant diversification.

Although the presence of additional lineage-specific responses cannot be excluded, our results suggest that an important component of the build-up of the pollination syndromes involves the convergent evolution of the expression across several hundreds of genes. These findings, combined with the transcriptomic resource that we gathered, open promising opportunities to extend our understanding of the evolution of the Gesneriaceae family in a genetic, ecological and evolutionary context. On a larger scale, the genes that we characterized should be further tested on model systems to clearly identify their functions and in other groups showing contrasting pollination syndromes to integrate and complement our knowledge on the key genes for floral transitions and the pace of floral evolution.

## Material and Methods

### Species selection, sample collection and RNA sequencing

We selected in the Gesneriaceae family three pairs of related species, each with contrasting functional group of pollinators (i.e., hummingbirds vs. bees; Table S1, Fig. 1A). Pollinator information for each species derived from field observations. Bee pollination was recorded for *Nemathanthus albus* (Wolowski et al., com. Pers.), *Paliavana tenuiflora* (Ferreira & Viana, 2010), *Sinningia eumorpha* (Sanmartin-Gajardo & Sazima, 2004), whereas hummingbird pollination was recorded for *N. fritchii* (Franco & Buzato, 1992), *Vanhouttea calcarata* (Sanmartin-Gajardo & Sazima, 2005), and *S. magnifica* (Vasconcelos & Lombardi, 2000, 2001). To verify the validity of pollination syndromes, we quantified flower morphologies using 10 floral traits available from the literature (Perret et al., 2007; Serrano-Serrano et al., 2015, Table S2). Floral morphospace was visualized using a Principal Component Analysis (PCA) as implemented in the R function *prcomp* (R Core Team, 2017).

Transcriptomic data was already available for two of the selected species (S. *eumorpha* and *S. magnifica;* Serrano-Serrano et al., 2017b). For the other species, the sample collection, RNA extraction, library construction and sequencing was performed as in Serrano-Serrano et al. (2017b; see also SI S1).

### Transcriptomes assembly and orthology inference

The transcriptome assembly, annotation and quality assessment for each species were performed as described in Serrano-Serrano et al. (2017b; see also SI S1). Orthologous groups (OGs) between the six species were inferred with OrthoMCL (v.2.0.9; Li et al., 2003), using the predicted proteins as input. OGs were classified according to the number of species and number of sequences composing them (Table S5, Figure S3). We were here interested in investigating changes in gene expression potentially associated with the convergent evolution of pollination syndromes. Our analyses thus mainly focused on OGs including sets of species that enabled us to contrast the two pollination syndromes. This requirement was met by OGs composed of at least the three species with the same pollination syndromes, resulting in the analysis for differential expression linked with pollination syndromes of 14,059 OGs. These OGs were composed by 3 (“pollination-syndorme-specific OGs”), 4 (“four-species OGs”), 5 (“five-species OG”) or 6 species (1-to-1 OG). For all categories, only OGs with a single expressed copy per species were considered. Additional information on the selection of OGs is available in SI S2.

### Raw and normalized gene expression, and overall comparison of the expression profile of the sequenced libraries

The raw expression of each gene in each tissue and species was obtained as described in SI S3. Briefly, we mapped RNA-Seq reads of each library to the assembled transcriptome of the corresponding species with Bowtie (Langmead et al., 2009). We estimated the raw expression of genes in each tissue and species with the *align_and_estimate_abundance.pl* script (Trinity tools, Haas et al., 2013; RSEM quantification metho, Li & Dewey, 2011).

Gene expression normalization was performed to account for differences in sequencing depth and in gene length within libraries and between species, and to correct for mean-variance dependency (Zwiener et al., 2014). The *rpkm* and *calcNormFactors* functions were used (edgeR package; Robinson et al., 2010) to account for differences in gene length and sequencing depth, respectively. The *voom* function in the limma R package (Ritchie et al., 2015, normalization method: *cyclic-loess*) was used to calculate the variance weights and to transform the normalized expression data ready for linear modeling. We accounted for replicated samples within conditions, batch effect and phylogenetic relationships by using an appropriate design matrix. Additional information is available in SI S4.

To investigate the overall quality and differentiation between RNA-Seq libraries, we considered the normalized gene expression matrix for 1-to-1 OGs and we compared the overall expression profile of floral and leaf tissues in the different species based on PCA and k-mean clustering as implemented in the factoextra R package (Kassambara & Mundt, 2015).

### Differential gene expression analysis between pollination syndromes

For 1-to-1 OGs, two different approaches were employed to investigate the genes differentially expressed between bee- and hummingbird-pollinated species. The first (i.e. “pairwise” differential expression analysis) consisted in first comparing the expression within the three pairs of related species differing in their pollination syndromes, and then investigating the overlap of the DEGs between the three species pairs. In the second approach (i.e. “global” differential expression analysis), we directly contrasted all bee-pollinated species against all hummingbird-pollinated species while accounting for phylogenetic relationship, thus considering species with the same pollination syndromes as replicates. While both approaches can detect the clearest cases of concerted changes in expression linked with the pollination syndromes (such as in Figure S6A-B), for less obvious cases the two approaches are complementary. For additional information, see SI S5 and Figure S6.

In the “pairwise” analysis, we first retrieved the normalized expression ready for linear modeling of the 1-to-1 OGs (see above). For each pair of related species, we fitted a gene-wise linear model and contrasted the gene expression of the related species in the different floral developmental stages while accounting for replicated samples within conditions and batch effect. For this, we used the *lmFit, contrasts.fit* and *eBayes* functions in limma R package (Ritchie et al., 2015). For each floral developmental stage within each pair of species, we identified significantly differentially expressed genes (DEG) with the function *decideTests* (method: *hierarchical*, p-value adjust method: *fdr).* An adjusted p-value threshold of 0.05 and minimum log2-fold change of log2(1.5) was chosen for a gene to be a DEG. The log2-fold change threshold corresponds to a difference in expression of at least 50% between the conditions. We then investigated the overlap of DEGs between species pairs for each floral developmental stages, resulting in DEGs specific to only one species-pairs, DEGs found in two species pairs, and DEGs shared by the three species pairs. DEGs that were found shared between species pairs but showing contrasting expression patterns (such as in Figure S6E) were not considered. We estimated the random expectation of the overlap of DEG between species pairs by randomly sampling a number of genes equal to those obtained in the pairwise analysis and distributing them in the 1 and −1 categories (where −1 and 1 respectively correspond to higher expression in bee- and hummingbird-pollinated species). We investigated the overlap of the randomly selected genes in each pairs of species. We repeated this 1,000 times to obtain the distribution of expected genes overlap between species pairs for each developmental stage. Additional information is reported in SI S8.

The “global differential expression analysis” was performed similarly to the “pairwise” one. We fitted the gene-wise model and we contrasted the gene expression of bee- and hummingbird-pollinated species in the different floral developmental stages. For each floral developmental stage we identified DEGs with the function *decideTests* (method: *hierarchical*, p-value adjust method: *fdr).* An adjusted -p-value threshold of 0.05 and minimum log2-fold change of log2(2) was required for a gene to be a DEG (see SI S5).

For OGs specifically found only in one of the pollination-syndromes, we tested if the gene expression was significantly higher than zero. For both “bee”- and “hummingbird”-specific OG, we first retrieved the normalized expression and performed one-sample t-test for each gene in each species and developmental stage while correcting for multiple-testing (*p.adjust* function in R, method *fdr*, threshold of 0.05). We then kept only genes showing expression significantly different from zero in the three species. Additional information on the approach used is provided in SI S6.

For four- and five-species specific OGs, we investigated whether the expression in all species with the same pollination syndromes was significantly higher than the expression in the species with contrasting phenotypes. For the species with missing expression levels, we assumed that their expression level was zero. We retrieved the normalized expression of the these OGs and for each developmental stage we contrasted the expression of the closely-related species using a two-sample t-test. Multuple testing correction was performed as described above (*p.adjust* function in R; method *fdr*, threshold of 0.05). We considered only OGs showing significant and concordant differential expression (i.e. same direction for the difference). Additional information is provided in SI S7.

### STRING, Gene Ontology (GO) and KEGG enrichment analysis

We considered all the genes with an expression difference associated with one or the other pollination syndromes and retrieved their annotation based on *Arabidopsis thaliana* genes IDs. We generated the protein-protein interaction network using STRING (v.11, Szklarczyk et al., 2014), using the *A. thaliana* genome as reference background (total of 27,413 distinct protein-coding genes considered as gene “universe”). Functional enrichment analyses (Gene Ontology and KEGGs) were performed within STRING, considering the categories to be enriched when the reported false discovery rate (fdr) was lower than 0.01. GO enrichment analysis was also performed with TopGO R package (*classic* algorithm, *fisher* statistic, p-value threshold of 0.05; Alexa & Rahnenfuhrer, 2016), using the whole dataset of tested genes (14,059 OGs) as the gene universe. The results that we obtained were similar to those obtained with the STRING GO enrichment analysis and are not reported. For additional information, see SI S9.

## Author contributions

MLSS, MP, and NS conceived the study, MLSS and AM contributed to the laboratory work and analysis, MLSS, AM, MP, and NS led the writing.

## Acknowledgments

We thank Andrea Komljenovic, Julien Roux, and Charlotte Soneson for helpful discussions and assistance. We thank Dessislava Savona Bianchi for the support in the laboratory, and the team of the Lausanne Genomic Technologies Facility (GTF) at the Center for Integrative Genomics (University of Lausanne), for the technology platform assistance. Computational resources were provided by the Vital-IT infrastructure of the Swiss Institute of Bioinformatics. We thank Alain Chautems for sharing his expertise about the species and the staff of the Jardin botanique de la Ville de Genève for the propagation and maintenance of the living collection of Gesneriaceae. Collection permits were granted by the Conselho Nacional de Desenvolvimento Científico e Tecnológico (CNPq) of Brazil (EX-15/89 and CMC 038/03). Images in figure 1C and 1D were kindly provided by Ivonne SanMartin-Gajardo. This work was supported by a PhD fellowship from the Faculty of Biology and Medicine to MLSS, the grant CRSII3 147630 from the Swiss National Science Foundation to NS, the grant 31003A_175655 from the Swiss National Science Foundation to MP, and from funding of the University of Lausanne.

